# In vivo Modulation of Intraocular and Intracranial Pressures Causes Nonlinear and Non-monotonic Deformations of the Lamina Cribrosa and Scleral Canal

**DOI:** 10.1101/2023.01.29.526113

**Authors:** Ziyi Zhu, Susannah Waxman, Bo Wang, Jacob Wallace, Samantha E. Schmitt, Elizabeth Tyler-Kabara, Hiroshi Ishikawa, Joel S. Schuman, Matthew A. Smith, Gadi Wollstein, Ian A. Sigal

## Abstract

**Purpose:** To evaluate changes in monkey optic nerve head (ONH) morphology under acutely controlled intraocular pressure (IOP) and intracranial pressure (ICP).

**Methods:** Seven ONHs from six monkeys were imaged via optical coherence tomography while IOP and ICP were maintained at one of 16 conditions. These conditions were defined by 4 levels for each pressure: low, baseline, high and very high. Images were processed to determine scleral canal area, aspect ratio, and planarity and anterior lamina cribrosa (ALC) shape index and curvature. Linear mixed effect models were utilized to investigate the effects of IOP, ICP and their interactions on ONH morphological features. The IOP-ICP interaction model was compared with one based on translaminar pressure difference (TLPD).

**Results:** We observed complex, eye-specific, non-linear patterns of ONH morphological changes with changes in IOP and ICP. For all ONH morphological features, linear mixed effects models demonstrated significant interactions between IOP and ICP that were unaccounted for by TLPD. Interactions indicate that the effects of IOP and ICP depend on the other pressure. The IOP-ICP interaction model was a higher quality predictor of ONH features than a TLPD model.

**Conclusions:** In vivo modulation of IOP and ICP causes nonlinear and non-monotonic changes in monkey ONH morphology that depend on both pressures and is not accounted for by a simplistic TLPD. These results support and extend prior findings.

**Translational Relevance**: A better understanding of ICP’s influence on the effects of IOP can help inform the highly variable presentations of glaucoma and effective treatment strategies.

## Introduction

Glaucoma is a progressive, irreversible optic neuropathy and the second-leading cause of vision loss in the world.^1,2^ The most prominent initial site of injury is the optic nerve head (ONH).^3–6^ Although the exact mechanism of neural tissue loss in glaucoma remains unknown, several studies suggest that mechanical insult contributes to the damage of neural tissues in glaucoma. Currently, the only modifiable risk factor for glaucoma is elevated intraocular pressure (IOP).

Elevated IOP is associated with an increased risk of mechanical insult at the ONH. This IOP-mediated mechanical insult can initiate the neurodegeneration characteristic of the disease. However, it is well established that there is marked variability between subjects in the sensitivity to elevated IOP. The fact that there exist large variations in sensitivity to IOP among individuals remains unexplained. Recent animal studies suggest that the intracranial pressure (ICP) could be the missing piece of the puzzle needed to explain the variation in sensitivity to elevated IOP.^7–10^

There are a few studies that measure the effect of IOP and ICP manipulation in animal models, including a canine model^11^ and a rat model.^12^ Morgan et al.’s study on a canine model suggested that the optic disc surface moved posteriorly under elevated IOP and moved anteriorly under elevated ICP. These changes were estimated based on assuming that the optic disc surface is a surrogate for the deeper tissues, which is now known not to reflect the true tissue mechanics.^13–15^ Zhao et al.’s work on a rat model also found that increased ICP could relieve the effect of elevated IOP on the retina and ONH structures. The authors of these studies interpret the results as suggesting that IOP and ICP could have counteracting effects such that an increase in one pressure could potentially cancel out the effect of the other being elevated.

Our group recently utilized a nonhuman primate model to quantify the acute effects of IOP and ICP, each at physiologic and pathologically elevated levels, on the ONH.^16^ Specifically, our previous study demonstrated the substantial effects of ICP on the scleral canal’s area, aspect ratio, and planarity, in addition to the anterior lamina cribrosa’s (ALC’s) depth and visibility. We also detected interactions between effects of ICP and those from IOP. ICP was found to affect the ONH’s sensitivity to IOP, thus potentially affecting susceptibility to glaucoma. In this previous work, we evaluated the effects on the ONH of IOP, ICP, and their interaction in a focused study but with limited statistical power. Specifically, we compared the effect of IOP at low and high ICP as well as the effect on ICP at low and high IOP. The study incorporated 4 eyes and only analyzed binary pressure conditions, i.e. low and high. This previous preliminary study evidenced the existence of IOP-ICP interaction. Computational modeling studies from us^17^ and others^18^ suggest that there are rich and complex effects of both IOP and ICP on the ONH, including nonlinear responses. Such effects cannot be determined from binary pressure conditions.

Following up the previous study, the present work aims to measure the effects of acute IOP/ICP changes more comprehensively on the ONH of monkey models. Our goal was to analyze how the effect of IOP/ICP changes as ICP/IOP changes in more subjects and pressure levels instead of the binary tests performed in the previous study. Specifically, we quantified deformations of the scleral canal and ALC under 16 acute, controlled combinations of IOP and ICP (low, baseline, high, and very high) in 7 eyes. ONH deformations were quantified using 3 scleral canal parameters (canal area, aspect ratio, and planarity) and 2 ALC intrinsic shape parameters (shape index (SI) and curvedness). This increased level of detail can allow us to better capture potential factor influences, including what may be crucial nonlinearities.

## Methods

We utilized rhesus macaque monkeys as a model for our in vivo experiments. Surgical procedures, pressure control and imaging procedures were as described before.^16^ For clarity we will describe the key elements here. Both IOP and ICP were independently controlled to allow for simultaneous, acute manipulation. Under 16 distinct pressure conditions, 7 eyes from 6 monkeys were imaged, in vivo, with optical coherence tomography (OCT). Each OCT image volume was processed into virtual radial slices on which ONH structures were manually delineated. Image processing was completed in FIJI.^19^ Custom scripts were used to compute scleral canal area, aspect ratio, and planarity as well as ALC shape index (SI) and curvedness from the delineations. Finally, we applied a linear mixed effects model to analyze the effect of IOP, ICP, and their interaction on ONH morphology.

### Animal Handling

All animal procedures followed the National Institute of Health (NIH) Guide for the Care and Use of Laboratory Animals, adhered to the Association of Research in Vision and Ophthalmology (ARVO) statement for the Use of Animals in Ophthalmic and Vision Research, and were in accord with a protocol approved by the Institutional Animal Care and Use Committee (IACUC) of the University of Pittsburgh. Before the experiment, a clinical examination was conducted to exclude eyes with gross abnormality. Each of 6 monkeys were prepared for imaging as described previously ^16^ Animals were initially sedated with 20 mg/kg ketamine, 1 mg/kg diazepam, and 0.04 mg/kg atropine. They were maintained on 1-3% isoflurane for the remainder of the experiment. Animals were put on a ventilator and given vecuronium bromide, a paralytic, intravenously at 0.04-0.1 mg/kg/hour to reduce drift in eye position throughout the experiment. Pupils were dilated using tropicamide ophthalmic solution 0.5% (Bausch & Lomb, Rochester, NY). Eyes were scanned while animals were in the prone position, with the head held upright and facing the OCT device. The corneal surface of each eye was covered with a rigid, gas permeable contact lens (Boston EO, Boston, MA) to preserve corneal hydration and improve image quality. The eyes were kept open using a wire speculum and the corneas were hydrated with saline between scans. The animals’ blood pressures and heart rates were monitored throughout the study.

### Manipulation of Intraocular and Intracranial Pressure

To control IOP, a 27-gauge needle was inserted into the anterior chamber and connected to a saline reservoir (**Figure 1a**). To control ICP, a lumbar drain catheter was flushed of air, inserted 2.5 cm into the lateral ventricle of the brain, and then connected to a saline reservoir. The IOP was determined by the height of the reservoir. The ICP was determined by a pressure recorder placed into the lateral ventricle, at least 5 mm away from the catheter (Codman ICP Express, Johnson & Johnson, Raynham, MA). Before using the pressure transducer, it was calibrated while submerged in saline solution. IOP and ICP values were controlled within 1 mmHg. This study included 16 pressure conditions (**Figure 1b**). The 16 conditions consisted of combinations of 4 levels of IOP and ICP: low, baseline, high, and very high. Examples include low IOP and low ICP, baseline IOP and low ICP, high IOP and low ICP, etc. The approximate respective mmHg values for each pressure group were 5, 15, 30, and 45mmHg for IOP and 5, 10, 25, and 35mmHg for ICP.^20^

**Figure 1.**
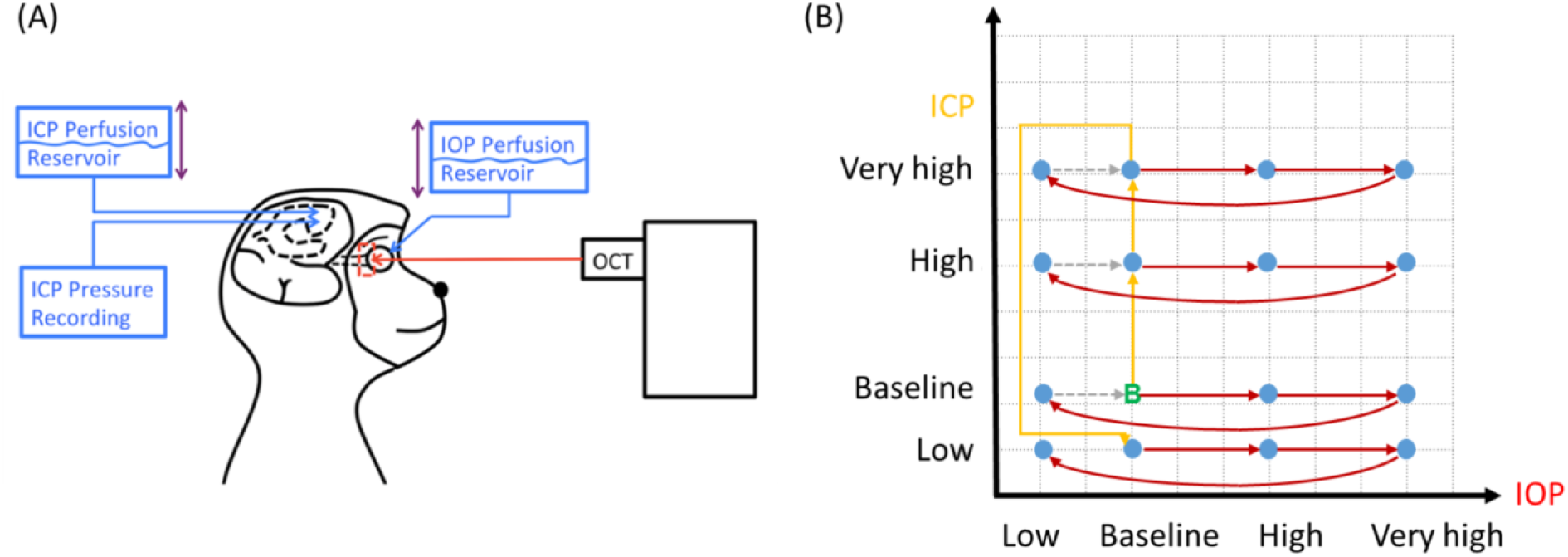
In vivo experiment set-up. (A) Monkey eyes were imaged with OCT while IOP and ICP were controlled using saline reservoirs. (B) Diagram of pressurization with imaging points (blue), started at baseline IOP and ICP (green). Between each set of IOP elevations (red arrows), ICP was changed stepwise from baseline to very high then back to low level (yellow arrows).

### Imaging

Monkey eyes were imaged using spectral domain optical coherence tomography (SD-OCT, Bioptigen, Research Triangle, NC) with a scan rate of 20,000 A-scans/second, modified with a broadband superluminescent diode (Superlum, Dublin, Ireland, λ = 870 nm, Δλ = 200 nm). OCT scans were centered on the ONH region (**Figure 1a**) with a size of either 3×3×2 mm or 5×5×2 mm and 512×512×1024 pixels sampling. Under each pressure condition, multiple scans were taken and scans with the best quality were used to perform manual delineation. Image quality criteria are detailed elsewhere.^21^ After each pressure manipulation, a minimum wait time of 5 minutes was observed before imaging to ensure that viscoelastic effects had dissipated. In addition, at each pressure we spent 20-30 minutes adjusting equipment and conducting the imaging. Image quality tended to decrease with increasing anesthesia time. To ensure that image quality remained high, we imaged only one eye from most animals (5 out of 6). All scans were re-sampled at 1 × 1 × 1 scale for analysis.^22^ Eyes vary in optical power and OCT systems are optimized for imaging human eyes. Hence, OCT images of monkey ONHs must be rescaled in the transverse dimensions. To set the dimensions, we followed the process described previously.^21^ Briefly, after the experiment, eyes were enucleated, processed for histology, and sections were imaged with polarized light microscopy. The images were reconstructed into 3D stacks and used to obtain eye-specific transverse scaling factors based on the dimensions of the scleral canal at BMO. Elsewhere we have shown that histological processing does not alter the scale of eye tissues.^21,23^ The determined scaling factors were then applied to the OCT images before the following analysis.

### Image Processing and delineation

We identified motion artifacts due to breathing and heartbeat as periodic patterns in the smooth structure of the Bruch’s membrane in the OCT slow scan direction. We mitigated these artifacts by translating individual B-scan images in the anterior-posterior direction. Radial re-slice was then performed on the OCT image volume using previously developed scripts in FIJI.^21^ Preliminary markings were made on the Bruch’s membrane opening (BMO) in the en face view prior to the re-slice process to determine the center of re-slicing (**Figure 2a**). Through the re-slice process, we obtained 18 virtual, radial slices for each image volume (**Figure 2b**). A Gaussian filter was applied to the radial image stack before delineation of ONH features to remove the background noise and improve image quality.

**Figure 2.**
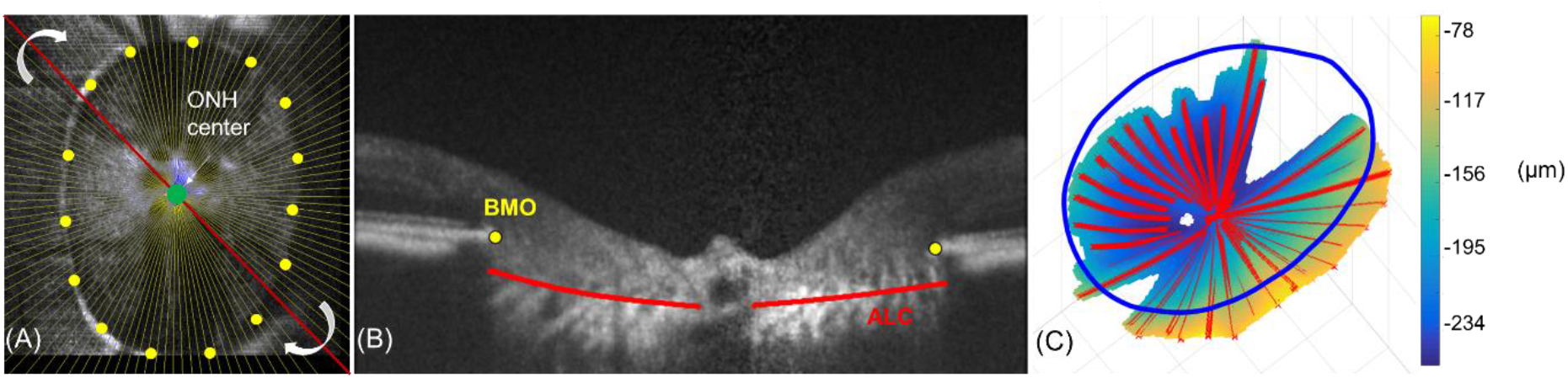
Example of our image analysis. (A) En-face view of BMO outline (yellow dots) used to determine ONH center (green) for virtual radial reslicing (red line). Motion artifacts in the slow scan direction were removed, and virtual radial slices are generated, centered at the centroid (green) of the scleral canal (B) Example markings of the BMO (yellow) and ALC boundary (red) on a virtual radial slice. Radial slices are then delineated for the anterior lamina cribrosa (red) and the scleral canal (yellow), measured at the Bruch membrane opening. (C) Example heat map of ALC depth with radial markings (red) and outline of best-fit BMO plane (blue).

Delineation of BMO and ALC in these radial slices was performed in FIJI by an experienced observer masked to both IOP and ICP conditions. The delineation process yielded 3D markings of these ONH structures (**Figure 2c**) which are anatomical landmarks often used in studies^24^ of ONH biomechanics using OCT.

### 3D reconstruction & registration

Delineations were imported with our custom scripts from FIJI to MATLAB, where 3D ALC surfaces were reconstructed using scattered data interpolation. A confidence map was also imposed to only regions with high reliability. The scleral canal, defined as the best-fit plane of the BMO, was used as a reference plane to calculate ALC depth. For each individual eye, the reconstructed ALC surfaces from scans of different pressure conditions were registered by aligning the center and principal axes of the scleral canal.

### Scleral Canal Area, Aspect Ratio, and Planarity

Scleral canal area was computed as the BMO’s projected area onto its best-fit plane. Canal aspect ratios were defined as the ratio between the major and minor principal axes (**Figure 2**). Planarity was computed as the mean distance from the BMO to the best-fit plane. Notice that, by definition, a planarity value of zero represents a flat plane and higher planarity values imply larger deviation from a flat plane.^6,25^

### Anterior Lamina Cribrosa: Shape Index and Curvature

The lamina cribrosa global shape index (SI, **Figure 3**) introduced by Thakku et al,^26^ is a novel parameter to characterize the shape of the lamina cribrosa. In brief, it was computed through the following steps. First, we created virtual radial sections of the ALC surface to obtain 180 arcs, each at 1 degree apart from its neighbors. We then computed the principal curvatures K1 and K2, which are the maximum and minimum curvature of the ALC surface among the 180 arcs (**Figure 3a**). Positive curvedness indicates a more posteriorly curved ALC while negative curvedness indicates a more anteriorly curved ALC. The SI and the curvedness (C) are given by the formulas provided in **Figure 3b**.

**Figure 3.**
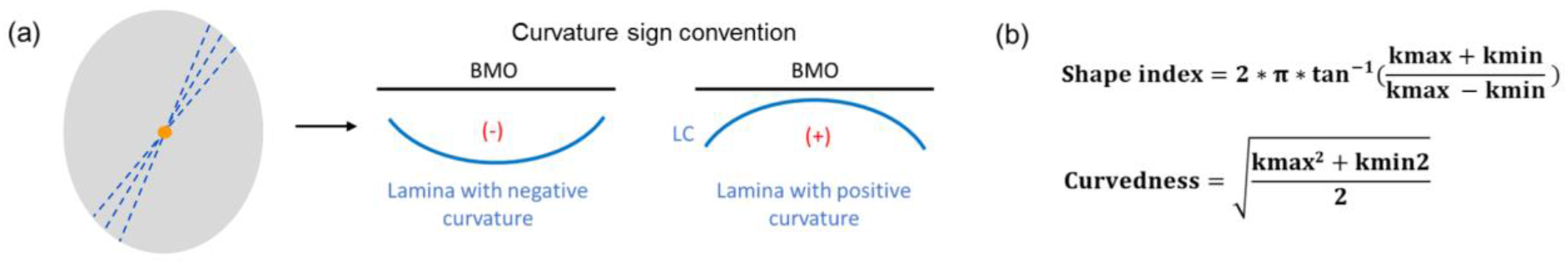
Computing lamina shape parameters. (a, left) Virtual radial slices (dashed blue lines) from each lamina surface (gray) centered at the centroid (orange point) of the scleral canal. (a, right) Arcs with negative curvature correspond to a concave ALC and positive curvature corresponds to a convex ALC. (b) Definitions of shape index and curvedness, calculated from the maximum and minimum principal curvatures of ALC surfaces (kmax, kmin).

### Data handling and Validation

When describing parameter changes, we defined positive changes as increase and negative changes as decrease, regardless of the parameter’s value being positive or negative. For example, a positive change of a negatively valued parameter still indicates an increment toward positive infinity.

To adapt to the relatively small subject population, we employed the method of bootstrapping to verify the stability and trustworthiness of the data, i.e. if the markings consistently reflected the behavior of the structures under pressure, as done previously.^16^ Within each test, 80% of the markings were randomly selected. These randomly sampled subsets were then used to construct the surface model and compute the described parameters. This procedure was repeated 10 times for each eye, generating 10 sets of bootstrapped results. For every parameter, the standard deviation across the 10 sets were computed. The values of the calculated parameters over the 10 random subsets were averaged to produce the bootstrapped results which were compared to the original results obtained using all data.

### Statistics

Linear mixed effects models were constructed to analyze the effect of IOP, ICP, and their interactions on each of the five parameters separately. IOP, ICP and their interaction were defined as fixed effects, where individual monkey and individual eye were defined as random effects. Note that the linearity in the models refers to each of the parameters, and that it is possible for the models to describe nonlinear responses if the nonlinearities are due to interactions. The input fixed effect data was centered to baseline pressure such that zero corresponded to 15 mmHg for IOP and 10 mmHg for ICP. A second set of models constructed using translaminar pressure difference (TLPD) as the only fixed effect were also tested. For both models, an alpha of 0.05 was used. The two models were then compared utilizing Akaike information criterion (AIC) to determine relative superiority. Following the same approach we have described elsewhere, we evaluated whether transforming the variables was necessary or helpful, for instance to satisfy statistical assumptions on the distribution of residuals, or allowed for better fits.^16,27^ We found that variable transformations were not helpful enough to compensate for the increased complexity and thus the results shown are untransformed.

## Results

### Data handling and bootstrapping validation

We aimed to image eyes at IOPs of 5, 15, 30, and 45mmHg and ICPs of 5, 10, 25, and 35mmHg. Imaging constraints resulted in small variations in pressures at the time of imaging. Ranges for the exact mmHg values for pressures in each pressure group (low, baseline, high, and very high) are detailed in **Supplementary Table 1**. All statistical tests were conducted with exact mmHg values.

The bootstrap test results indicated that manual marking quality was consistent such that partial omission of markings led to minimal changes in the outcome of calculated parameters (**Table 1**). The standard deviation of the results from the 10 random bootstrap subsets averaged to 8.75 × 10-3, 8.18 × 10-3 mm^2^, 6.09 × 10-1 um^2^, 2.05 × 10-2, and 1.230 × 10^−5^ mm^−1^ for aspect ratio, area, planarity, SI, and curvedness, respectively. This variation was minimal compared to the standard deviation of the parameters across different pressure conditions as presented above, showing high consistencies across the 10 randomly sampled bootstrap subsets. The mean absolute percentage difference between the bootstrap results and the original results were 0.41%, 0.15%, 9.77%, 3.30%, and 3.52% for the five parameters. The differences were all below 10%, suggesting that using a randomly sampled subset of the markings caused only limited alteration in the results.

**Table 1.** Bootstrap test results. Mean inter-subset standard deviation, mean % difference from original results, and the range of difference are included for each ONH feature analyzed.

| % | Aspect Ratio | Area (mm<sup>2</sup>) | Planarity (μm) | Shape Index | Curvedness (μm<sup>-1</sup>) |
| --- | --- | --- | --- | --- | --- |
| Mean Inter-subsets STD | 8.75 x 10<sup>-3</sup> | 8.18 x 10<sup>-3</sup> | 6.09 x 10<sup>-1</sup> | 2.05 x 10<sup>-2</sup> | 1.23 x 10<sup>-2</sup> |
| Mean % Difference (abs value) to original results | 4.10 x 10<sup>-1</sup> | 1.50 x 10<sup>-1</sup> | 9.77 | 3.30 | 3.52 |
| Range of Difference | -1.32 — 0.08 | -0.87 — 0.14 | -44.65 — 20.62 | -44.71 — 6.09 | -5.07 — 19.40 |

### Complex changes in ONH morphology with IOP and ICP

Average ± standard deviation values for scleral canal aspect ratio, area, planarity, and ALC SI and curvedness at baseline IOP and ICP were 1.38 ± 0.08, 2.24 ± 0.42 mm^2^, 8.56 ± 5.3 μm, −0.81 ± 0.07 and 0.33 ± 0.11 mm^−1^ respectively. With IOP and/or ICP adjustment, the change in these 5 parameters from baseline ranged respectively from −7.8% to +3.2%, −7.3% to +21.4%, −47.7% to +392.2%, −25.0% to +205%, and −85.3% to +27.8%. Here, the larger than 100% increase in ALC SI corresponds to ALC shape inversion, i.e. from cup to cap or vice versa, leading to the change in sign of SI from negative to positive.

Complex, non-linear patterns of ONH morphological changes with change in IOP and ICP were observed that could not be captured by consideration of TLPD alone. **Figures 4** and **5** demonstrate changes of all measured morphological parameters with changes in IOP and ICP in each eye. IOP and ICP data is displayed in binned groups of low, baseline, high, and very high pressures (**Figure 4**) and as raw mmHg values (**Figure 5**). Eye-specific differences in responses to IOP and ICP were observed. Interestingly, contralateral eyes (eyes 3 and 4) appeared to have lower variability between them in their responses in comparison to eyes from other monkeys. Averaged responses revealed potential trends. Percent change in area, SI, and curvedness demonstrated a low amount of variability between eyes while planarity demonstrated a considerably higher degree of variability. Changes in area and curvedness appeared to be inversely correlated, where increases in area corresponded with decreases in curvedness. The pressure condition under which most eyes demonstrated the greatest amount of morphological change was at low IOP and high/very high ICP. On average, area, planarity, and SI were found to increase under these conditions while aspect ratio and curvedness were found to decrease.

**Figure 4.**
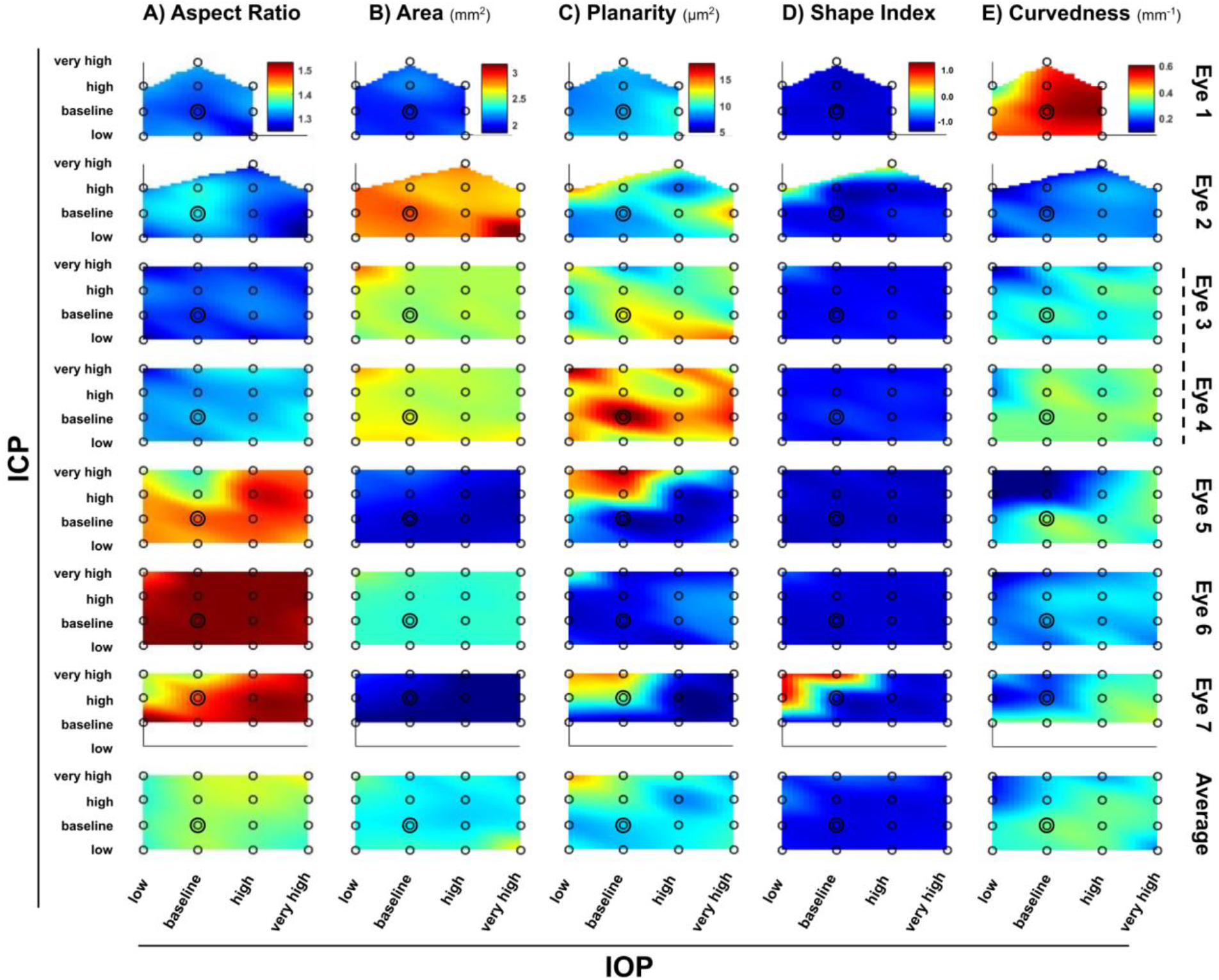
Colormaps demonstrating changes in (A) scleral canal aspect ratio, B) canal area, C) canal planarity, D) ALC shape index, and E) ALC curvedness with IOP and ICP. Red: increase, blue: decrease. Dashed line indicates contralateral eyes. IOP and ICP values were binned into low, baseline, high, and very high pressure groups according to the mmHg values in Supplementary Table 1.

**Figure 5.**
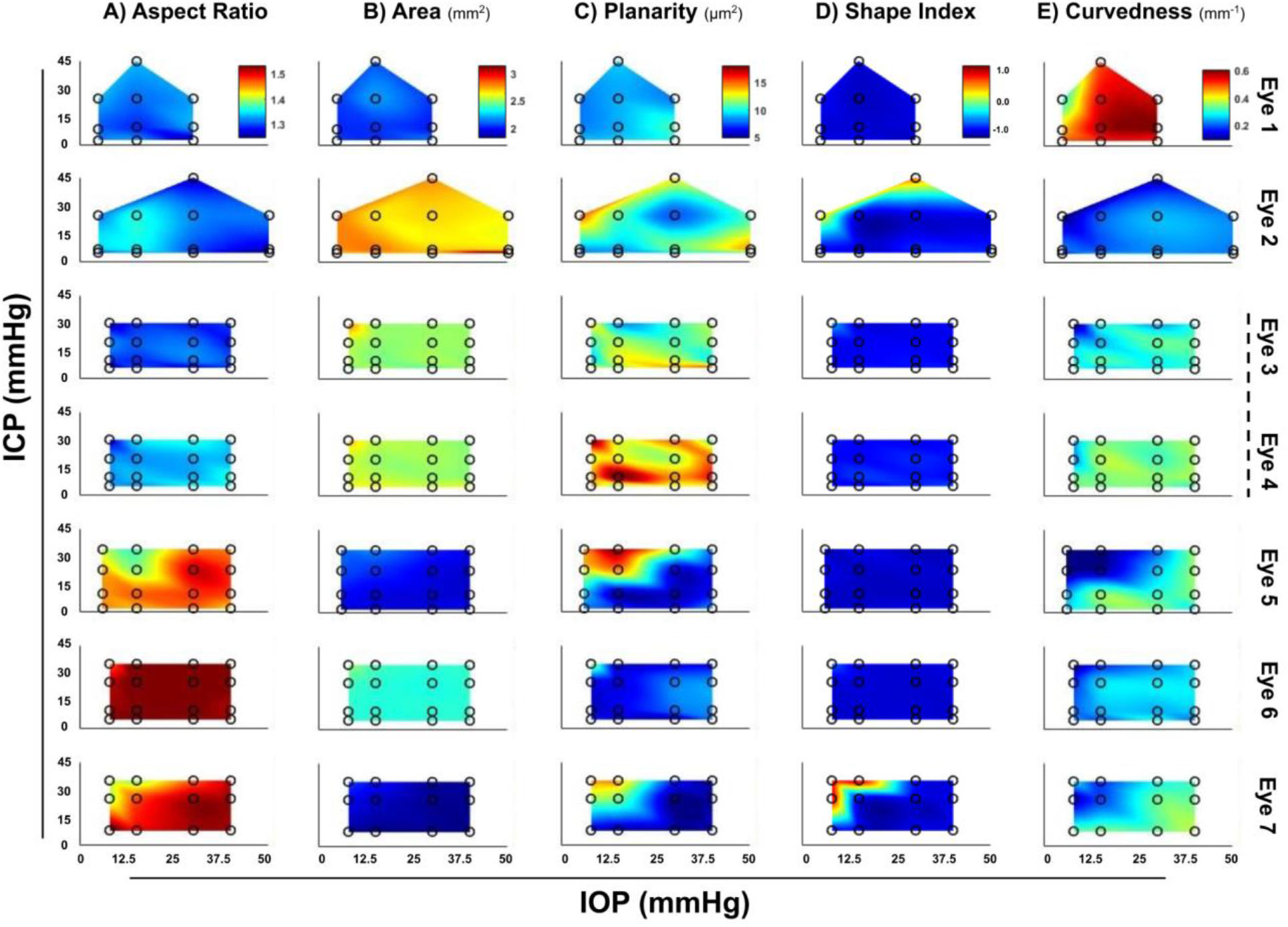
Colormaps demonstrating changes in (A) scleral canal aspect ratio, B) canal area, C) canal planarity, D) ALC shape index, and E) ALC curvedness with IOP and ICP. Red: increase, blue: decrease. Dashed line indicates contralateral eyes. IOP and ICP mmHg values are displayed without the binning used in Figure 4.

The SI distribution of monkey eyes versus human eyes is shown in **Figure 6**. The ALC SI of monkeys measured in this study was largely between −0.9 and −0.6, which corresponds to shapes between rut and cup. In human subjects, however, most SIs were distributed between −0.7 and 0.^26^ This difference was due to the absence of the central ridge, which forms a characteristic saddle shape in human ALC. Due to this difference, monkey ALC did not form a saddle shape, even under extreme pressure, but instead reversed its curvature and changed directly from a cup to cap shape.

**Figure 6.**
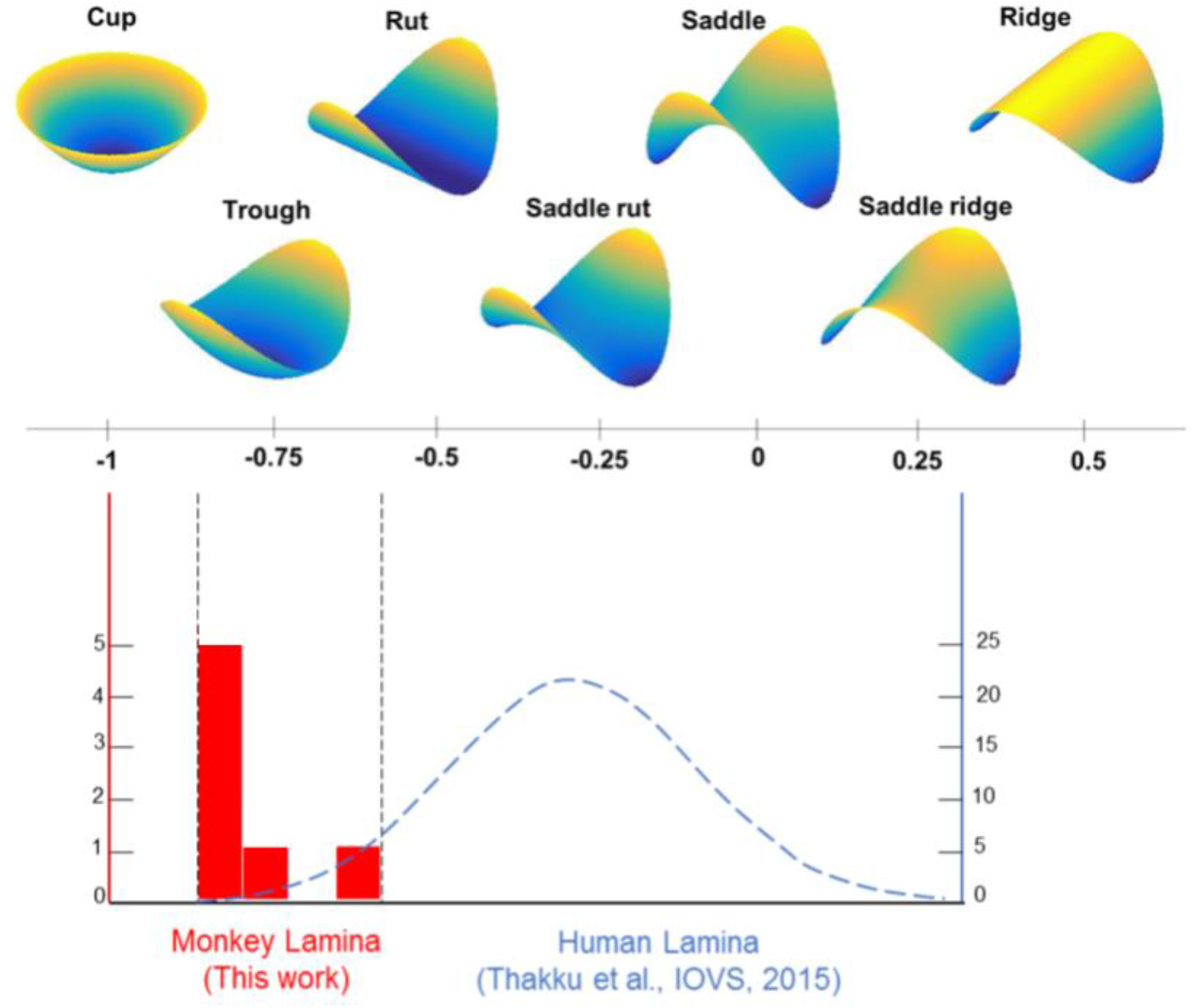
Comparing monkey and human ALC shape index (SI) at baseline IOP. Top panel is visualization of the characteristics that SI represents in different ranges. Bottom panel is the overlay of the distribution of human lamina SI in Thakku’s study and monkey SI in our study under baseline. The baseline monkey laminar shapes are mostly trough and are different from those of humans, which typically had saddle or rut shapes.

### Superior prediction of ONH morphology by IOP-ICP interaction than TLPD alone

Linear mixed effects models demonstrated significant interactions of IOP and ICP for all ONH morphological parameters. These parameters were scleral canal aspect ratio (p<0.001), canal area (p<0.001), canal planarity (p<0.001), ALC SI (p=0.012), and ALC curvedness (p<0.001). Fitted trend-lines for all parameters at different pressure conditions are shown in **Figure 7**. Corresponding p-values and coefficient summaries shown in **Table 2**.

**Table 2.**
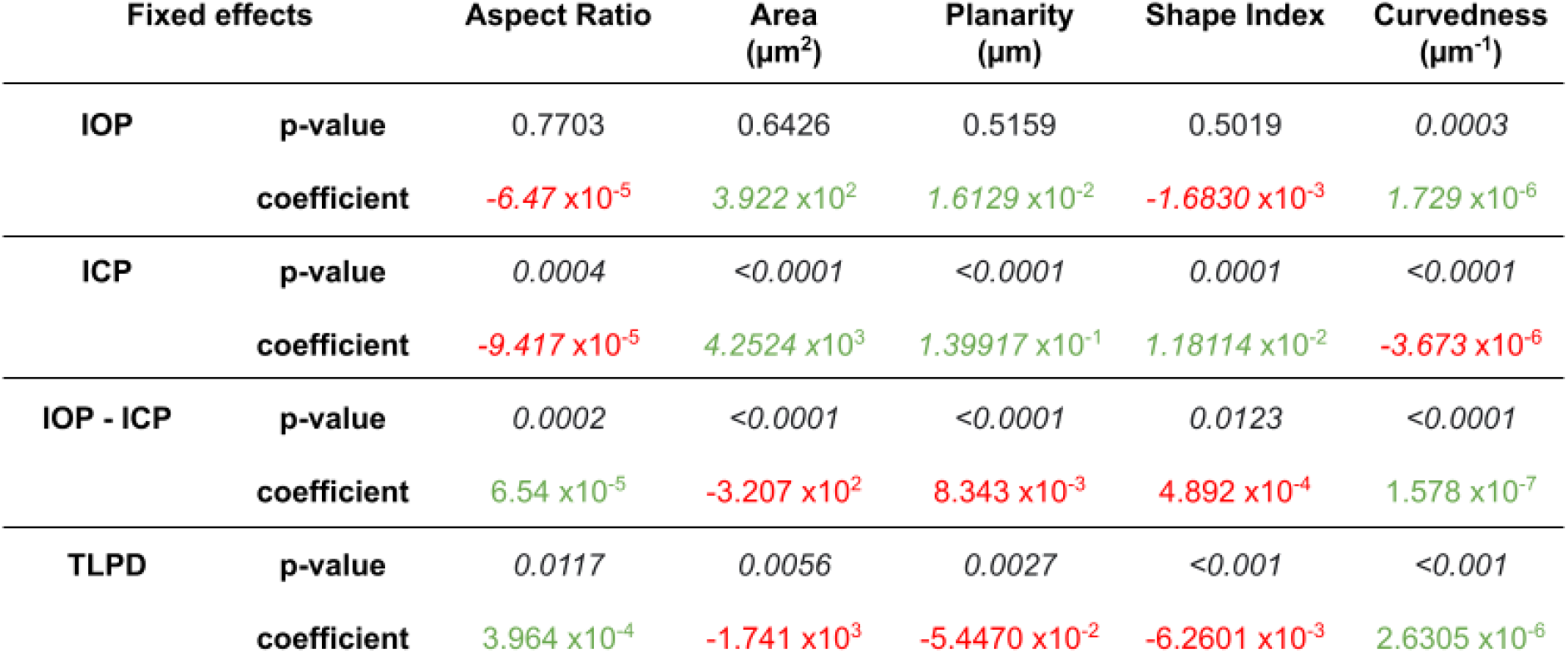
Linear mixed effects models p-values and coefficient summaries for models considering IOP, ICP, IOP-ICP interaction, and TLPD. Positive coefficient values are shown in green and negative coefficient values in red. The p-values below 0.05 are italicized.

| Fixed effects | | Aspect Ratio | Area<br>($\mu\text{m}^2$) | Planarity<br>($\mu\text{m}$) | Shape Index | Curvedness<br>($\mu\text{m}^{-1}$) |
| --- | --- | --- | --- | --- | --- | --- |
| IOP | p-value | 0.7703 | 0.6426 | 0.5159 | 0.5019 | 0.0003 |
|  | coefficient | -6.47 x10<sup>-5</sup> | 3.922 x10<sup>2</sup> | 1.6129 x10<sup>-2</sup> | -1.6830 x10<sup>-3</sup> | 1.729 x10<sup>-6</sup> |
| ICP | p-value | 0.0004 | <0.0001 | <0.0001 | 0.0001 | <0.0001 |
|  | coefficient | -9.417 x10<sup>-5</sup> | 4.2524 x10<sup>3</sup> | 1.39917 x10<sup>-1</sup> | 1.18114 x10<sup>-2</sup> | -3.673 x10<sup>-6</sup> |
| IOP - ICP | p-value | 0.0002 | <0.0001 | <0.0001 | 0.0123 | <0.0001 |
|  | coefficient | 6.54 x10<sup>-5</sup> | -3.207 x10<sup>2</sup> | 8.343 x10<sup>-3</sup> | 4.892 x10<sup>-4</sup> | 1.578 x10<sup>-7</sup> |
| TLPD | p-value | 0.0117 | 0.0056 | 0.0027 | <0.001 | <0.001 |
|  | coefficient | 3.964 x10<sup>-4</sup> | -1.741 x10<sup>3</sup> | -5.4470 x10<sup>-2</sup> | -6.2601 x10<sup>-3</sup> | 2.6305 x10<sup>-6</sup> |

**Figure 7.**
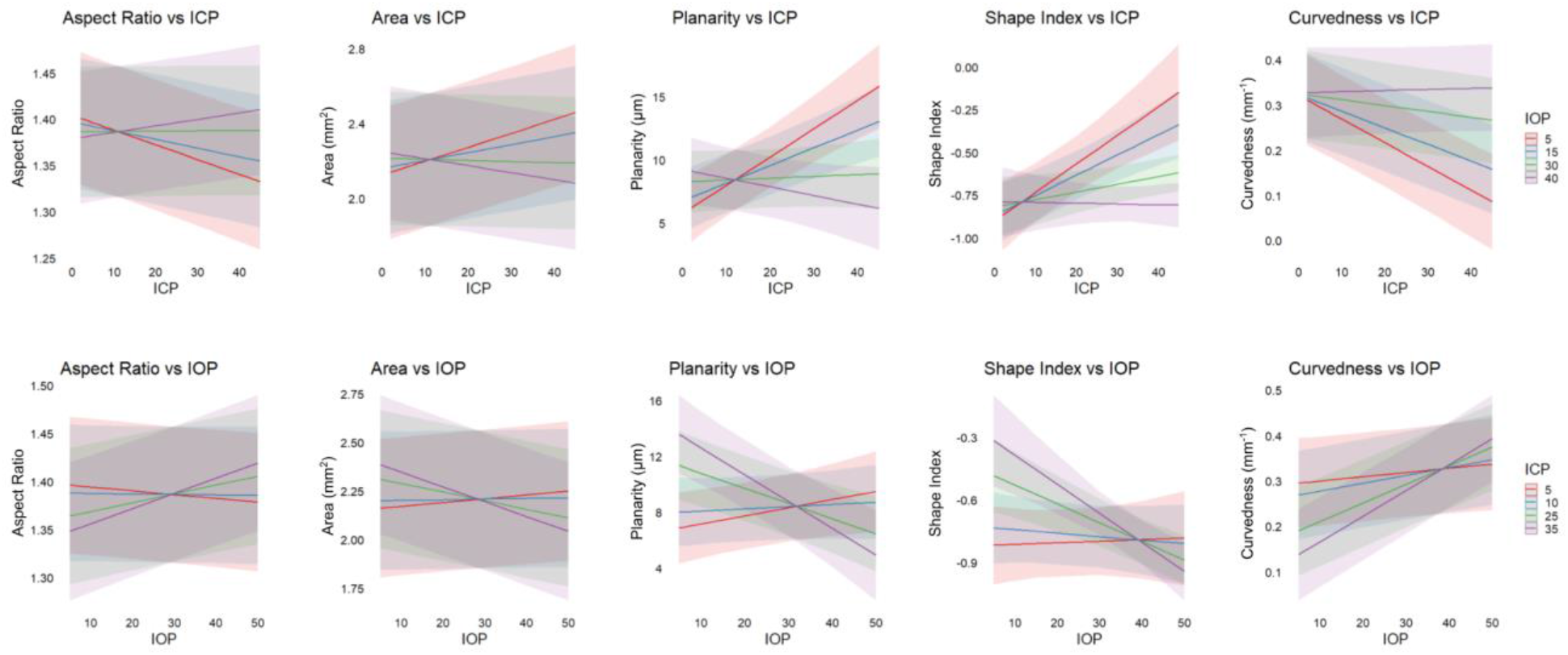
Linear mixed model results. Predicted effects of IOP-ICP combinations on ONH morphological parameters. Pressures: red = low, blue = baseline, green = high, purple = very high.

These significant interactions suggested that both the effect of IOP and ICP depend on the condition of the other pressure. In line with surface plots shown in **Figure 4 and 5**, the effect of IOP elevation on each morphological parameter was more prominent at elevated ICP. IOP elevation at high ICP considerably increased the scleral canal aspect ratio and decreased the canal area and planarity, leading to a more elliptical and flatter canal. Furthermore, IOP elevation at elevated ICP decreased SI and increased curvedness, resulting in a more posteriorly curved ALC. The effect of elevated ICP, on the other hand, was in general more prominent under baseline and low IOP. Increasing ICP led to a decrease in scleral canal aspect ratio and an increase in area and planarity, causing the canal to expand and become more circular and tilted. Increased ICP at low and baseline IOP also caused an increase in SI and a decrease in curvedness, anteriorly deforming the ALC to be less cup-like and less curved. Such effects, however, were substantially reduced or even reversed under elevated IOP.

TLPD similarly had significant effects on all five parameters: the scleral canal aspect ratio (p= 0.012), canal area (p=0.006), canal planarity (p=0.003), ALC SI (p<0.001), and ALC curvedness (p<0.001). The TLPD model is summarized in **Table 2**. We compared the AIC of the model accounting for IOP-ICP interaction and the model accounting for TLPD (**Table 3**). AIC was used as an indicator of prediction error and a metric of relative quality of each model. Lower AIC indicated a superior model. For all five parameters, the IOP-ICP interaction model, with a substantially smaller AIC, was of higher quality than the TLPD model.

**Table 3.** The AIC of the model accounting for IOP-ICP interaction and the model accounting for TLPD. For all five ONH parameters, the IOP-ICP interaction model, with a substantially smaller AIC, was of higher quality than the TLPD model.

| Parameters | AIC: IOP-ICP interaction model | AIC: TLPD model | Difference |
| --- | --- | --- | --- |
| Aspect Ratio | -408.5 | -398.2 | -10.3 |
| Area | -142.4 | -123.9 | -18.5 |
| Planarity | 509.7 | 524.9 | -15.2 |
| Shape Index | 48.4 | 51.8 | -3.4 |
| Curvedness | -269.9 | -254.8 | -15.1 |

## Discussion

In this study, we performed a comprehensive analysis on the effects of acute IOP and ICP changes, as well as the effects of their interaction, on ONH morphology. Our data was collected through experiments on 7 eyes from 6 monkeys. For each eye, we aimed to measure 5 morphological parameters at each IOP-ICP combination: canal aspect ratio, canal area, canal planarity, ALC SI, and ALC curvedness. We allowed 4 possible pressure levels for IOP and ICP low, baseline, high, and very high; and aimed to perform measurements for 16 conditions on each eye (combinations of the 4 pressure levels for IOP and ICP). For bootstrap analysis, randomly sampled markings were tested and shown capable of recovering the original results with small variations across the randomly sampled subsets. Bootstrapping tests suggested the results observed in the 6 healthy monkey subjects are reliable. Additional studies with a greater number of animals are necessary to characterize generalized effects of IOP/ICP on ONH responses at a population level. We are not aware of any other studies describing the effects of simultaneous IOP and ICP control on the monkey ONH at this level of detail.

Our study demonstrated three key findings. 1) IOP-ICP interaction significantly affects deformation of ONH features, revealing a complex, non-linear relationship between the effects of IOP and ICP. 2) These relationships between IOP and ICP effects can inform conditions under which small variations in pressure can have large effects on the ONH and vice versa. This may help inform high and low risk conditions. 3) IOP and ICP considered together are better predictors of ONH deformation than TLPD alone. We expand upon each of these findings below.

### IOP-ICP interaction significantly affects ONH morphology

As a main modifiable risk factor in glaucoma, IOP^28^ is known for its potentially damaging biomechanical effects on the ONH. Recent evidence has suggested the potential effects of ICP^7–9,29^ and has raised interest in studying its role in the ONH deformation and neural tissue damage. This study is not the first to report such interaction but provides additional ONH measures and more detailed data from monkey models as supportive evidence. The observations in the present study concurred with previous results that ICP changes could lead to significant deformation of the canal and ALC. However, it is worth noting that in most cases, the impact of ICP is reduced when IOP is elevated and does not always cause substantial deformations. This may be accounted for through IOP-ICP interaction.

A primary finding of the current study was the significant effect of interaction between IOP and ICP on all 5 computed parameters, which reflect deformations in both the scleral canal and the ALC. The presence of an interaction between IOP and ICP indicates that the effects of IOP and ICP depend on the level of one another and should not be considered separately. In general, the effect of IOP was more prominent when ICP was high, while the effect of ICP was more prominent when IOP was low. In some cases, the dependence was large enough to reverse the direction of parameter changes (**Figures 4, 5, 7**). This is in contrast with assumptions that the effects of IOP and ICP are linearly correlated.

In a range of conditions, the interaction between IOP and ICP can be described as a counteracting effect in a sense that increasing one pressure will reduce or inverse the effect of the other pressure. Specifically, simultaneously increasing IOP and ICP led to less ONH morphological changes than elevating ICP alone. Consider, for example, a process that first increases ICP and then increases IOP. According to the interaction effects (**Figure 7**), when ICP is raised, we would expect to observe the canal expand and become more tilted while the ALC becomes more anteriorly curved when ICP is raised. Later, as IOP is increased, the opposite effects would take place, contracting and flattening of the canal and a posterior curving of ALC. However, if we consider the above process again but first increase IOP then ICP, we would observe minimal or moderate deformation in both the canal and ALC throughout the process. Thus, the counteracting effects between IOP and ICP observed in this study originated from the interaction between IOP and ICP and does not necessarily imply that IOP and ICP have the opposite effect. Acutely elevated IOP alone did not lead to substantial deformations in the ONH structures but instead played a role of suppressing the effect of ICP. The effects of IOP under baseline ICP were not significant (**Table 2**, top row). This is in contrast with the effects of ICP under normal IOP. Several things must be taken into consideration when interpreting this result.

First, in a model with interaction of terms, the coefficients and p-values of the main effects in **Table 2** only represent their effect when the rest of the main effects were zero, which in our case corresponds to baseline. Clearly, IOP’s lack of significance at baseline ICP does not necessarily mean it has no significant impact overall. The fact that the effect of IOP and ICP significantly depend on each other implied that IOP played a crucial role in determining ONH deformations. As a result of the interaction, IOP had a much stronger effect under high ICP than low ICP.

Second, the difference between acute and chronic pressure change should be considered. Our experiment measured responses under acute IOP and ICP manipulations and did not include any measure on the effects of long-term elevated pressures, which are presumably more influential. Effects of short-term IOP elevation under normal ICP could intrinsically be more subtle and harder to capture. There are previous studies that reported no significant correlation between LC displacement and acute IOP manipulation in both human and monkey models.^13,30,31^

Moreover, few studies have measured the effects of IOP under directly controlled or monitored normal ICP and no previous studies had addressed the change of ALC shape under simultaneously controlled IOP and ICP. Our results serve as a further confirmation of the observations from these studies under better controlled conditions. There were studies that explored the displacements^11^ or strains^32^ of the LC under monitored IOP and ICP but were not well suited for comparison here, considering the fundamental difference between SI and displacement or strains. Nevertheless, it is worth noting that the parameters measured in both studies changed more prominently at lower pressures, which was not clearly seen from **Figure 7** in the present study.

### Non-linear changes in ONH morphology with IOP and ICP may inform risk

Due to the presence of IOP-ICP interaction, there are certain ICP conditions at which small changes in IOP may most affect the ONH. Conversely, we observed ICP conditions at which large changes in IOP had minimal effects. For example, while baseline ICP resulted in low to moderate effects of IOP, elevated ICP greatly increased the effect of IOP. Because the effects of IOP are dependent upon ICP, this indicates ICP as an important experimental variable to consider in studies evaluating the effects of IOP. In this instance, it may account for the difference between an ONH that experiences minimal deformation and an ONH that experiences significant deformation.

Clinically, this points to ICP potentially being the difference between neuroprotection and progressive neuropathy at a given IOP. IOP-ICP interaction is a potential risk factor for vision loss. Currently, there are yet to be robust explanations for why some individuals with high IOP do not develop glaucoma, why some individuals with healthy IOP develop normal-tension glaucoma (NTG), and why some glaucoma patients with IOP maintained at a safe level continue to experience neuropathy. IOP-ICP interaction may provide some insight.

In clinical studies, ICP was found to be lower in patients with NTG. Interestingly, we observed limited effects of IOP at low ICPs, suggesting that further reduction of IOP in these cases would not be of substantial therapeutic benefit. Our data are in line with clinical outcomes in which IOP reduction in NTG cases led to limited neuroprotective effects.^33^ Interestingly, ICP was found to be significantly higher in patients with ocular hypertension but no signs of glaucoma, suggesting a protective effect of high ICP at high IOPs.^8,33^ In line with these findings, we similarly see small effects of high IOP on the ONH at high ICP.

Although IOP and ONH morphology are readily measured in the clinic, measurement of ICP is invasive and therefore poses unwarranted risks. IOP and ONH morphology examined together, however, may provide evidence of IOP-ICP conditions and allow for more informed treatments. For example, if a patient with healthy IOP exhibits optic neuropathy, greater than average scleral canal area, planarity, and ALC SI with reduced ALC curvedness and scleral canal aspect ratio, this may indicate that ICP is high. This may suggest that IOP reduction would be of limited benefit and may possibly even exacerbate deformations. Other therapies may be more effective. In fact, our data suggest, in this instance, that IOP increase could bring ONH morphology back to a healthier, baseline state. As our work was conducted in 7 eyes of 6 monkeys, further studies are needed to make generalized conclusions about which ONH morphological features may best inform decision-making with high confidence.

### IOP-ICP interaction is a better predictor of ONH deformation than TLPD

Using AIC as the metric for model suitability, we found that the IOP-ICP model, which accounts for interaction effects between pressures, had a superior performance to the model with TLPD as the only fixed effect.

Studies have found that IOP and ICP could produce opposite effects on the ONH,^11,12^ thus supporting the hypothesis that TLPD, instead of IOP, is a better measure to estimate the effect of pressures in the eye. Evidence was found by previous studies that TLPD played an important role in the deformation of the ONH.^8,11^ However, although TLPD is a parameter that accounts for the effect of both IOP and ICP, it assumes a simple linear relationship with only first order terms between IOP and ICP. It similarly assumes that the two pressures add up to one net pressure exerted. Models utilizing TLPD cannot account for second order terms such as possible interactions between IOP and ICP. In the present study, the effects of two pressures combined were more complex than the effects of TLPD alone. Reducing ICP and elevating IOP led to different effects, even when the resulting TLPD was equivalent. For example, the effects caused by lowering ICP under low or normal IOP were not the same as those caused by increasing IOP under low or normal ICP for most parameters, where the effect of the former was either insubstantial or opposite to the later.

TLPD has been explored as a predictor and risk factor for neural tissue damage and glaucoma.^8,11,12^ Some studies have found a significant correlation between higher-than-normal TLPD and glaucoma,^8^ while there was also a study that found evidence against the measure of TLPD.^34^ In the present study, our results showed that TLPD changes were significantly correlated with changes in both canal and ALC parameters, which supported the hypothesis that TLPD serves as an indicator of ONH deformations. Although the effect of TLPD was significant, it may lead to an over-simplified model, given that both IOP and ICP were known. While TLPD was found to be a parameter that provides important insights, we have shown that there can be more than a simple canceling effect between IOP and ICP. Evidence found in the present study suggests that a more robust practice would be to take into account both IOP and ICP. The two should be considered as interacting factors when examining the effect of pressures on the ONH.

In this study, 3D deformations of the ONH resulting from changes in both IOP and ICP were measured in vivo via OCT imaging. Imaging in vivo avoids artifacts that could arise as a result of histological processing.^35,36^ This work was a comprehensive study of 7 eyes from 6 monkeys, each under 16 pressure condition combinations. Few studies have focused on the effects of the interaction between IOP and ICP. There have been in-depth studies on the mechanical effect of IOP, but few of them were conducted under controlled or even monitored ICP. In this study, we obtained enough data to perform regression analysis and test the significance of the effects caused by ICP and its interaction with IOP, thus testing the results of our previous study on a larger data set.

Mechanical insult to the LC, where the RGC axon loss takes place, plays a crucial role in the cause of glaucoma.^37,38^ Many studies used ALC depth (usually measured with respect to the BMO plane)^13,24^ to characterize LC shape. However, it is possible for ALC to have different shapes when its mean depth stays the same. In Tun et al’s previous study, they reported this issue with evidence that significant shape changes occurred in the ALC even though no significant change in depth took place.^39^ Our study employed novel parameters including ALC SI and curvedness to better describe the change of shape of the ALC and understand its deformation under pressure.

Another advantage of the ALC SI is that it can be computed solely based on the ALC surface and does not depend on the BMO reference plane. Calculating depth requires a reference by its definition. Although BMO was considered as a relatively stable structure and has been commonly used as a reference in previous studies,^24,25^ it could still experience displacements under pressure. In this study, the scleral canal opening experienced significant deformation under pressure. It thus may potentially alter the position of the BMO plane relative to the ALC. Employing the ALC SI eliminates artifacts caused by possible movements of the BMO plane.

As human and monkey ALC bear some different characteristics, SI and curvedness results from monkey subjects need to be carefully interpreted. The SI distribution of monkey eyes versus human eyes is shown in **Figure 6**. The ALC SI of monkeys measured in this study was mostly within −0.6 and −0.9, which correspond to shapes between rut and cup. In human subjects, however, the SI was largely distributed between −0.7 and 0.^26^ The difference was due to the absence of the central ridge, which forms a characteristic saddle shape in human ALC. Due to this difference, monkey ALC did not form a saddle shape even under extreme pressure but instead reversed its curvature and changed directly from cup to cap. Determining exactly how the monkey ALC deformation behavior maps to human ALC deformations under the same conditions will require future studies.

Although the linear mixed effects models captured well the effects of IOP, ICP and their interactions, without the need for variable transformations, the nonlinear relationships between the parameters suggest that future studies may benefit from considering more complex models. This could be complicated because fitting nonlinear models accurately usually requires more experimental data. An alternative is to enrich the statistical model fitting using mechanistic relationships that can be derived from computational models.^17,18,27^

In conclusion, we aimed to explore patterns of interaction between IOP and ICP in a monkey model. We demonstrate that IOP-ICP interaction significantly affects ONH feature morphology. Importantly, non-linear relationships between IOP and ICP effects may help inform high and low risk pressure conditions. We observed conditions under which small variations in pressure had large effects on the ONH and, conversely, conditions under which large variations in pressure had small effects on ONH morphology. Despite the use of TLPD in studies of ONH deformation, the effect of IOP and ICP considered in tandem was found to be a superior predictor of ONH morphology than TLPD. These findings indicate the importance of considering IOP, ICP, and their interactions in studies of ONH biomechanics.

## Supplementary material

**Supplementary Table 1.** Ranges of mmHg values for low, baseline, high, and very high pressures of IOP and ICP.

| Pressure groups | IOP<br>(min-max mmHg) | ICP<br>(min-max mmHg) |
| --- | --- | --- |
| Low | 5-9 | 2-6 |
| Baseline | 10-20 | 7-16 |
| High | 21-30 | 17-29 |
| Very high | 31-50 | 30-45 |

**Supplementary Table 2.**
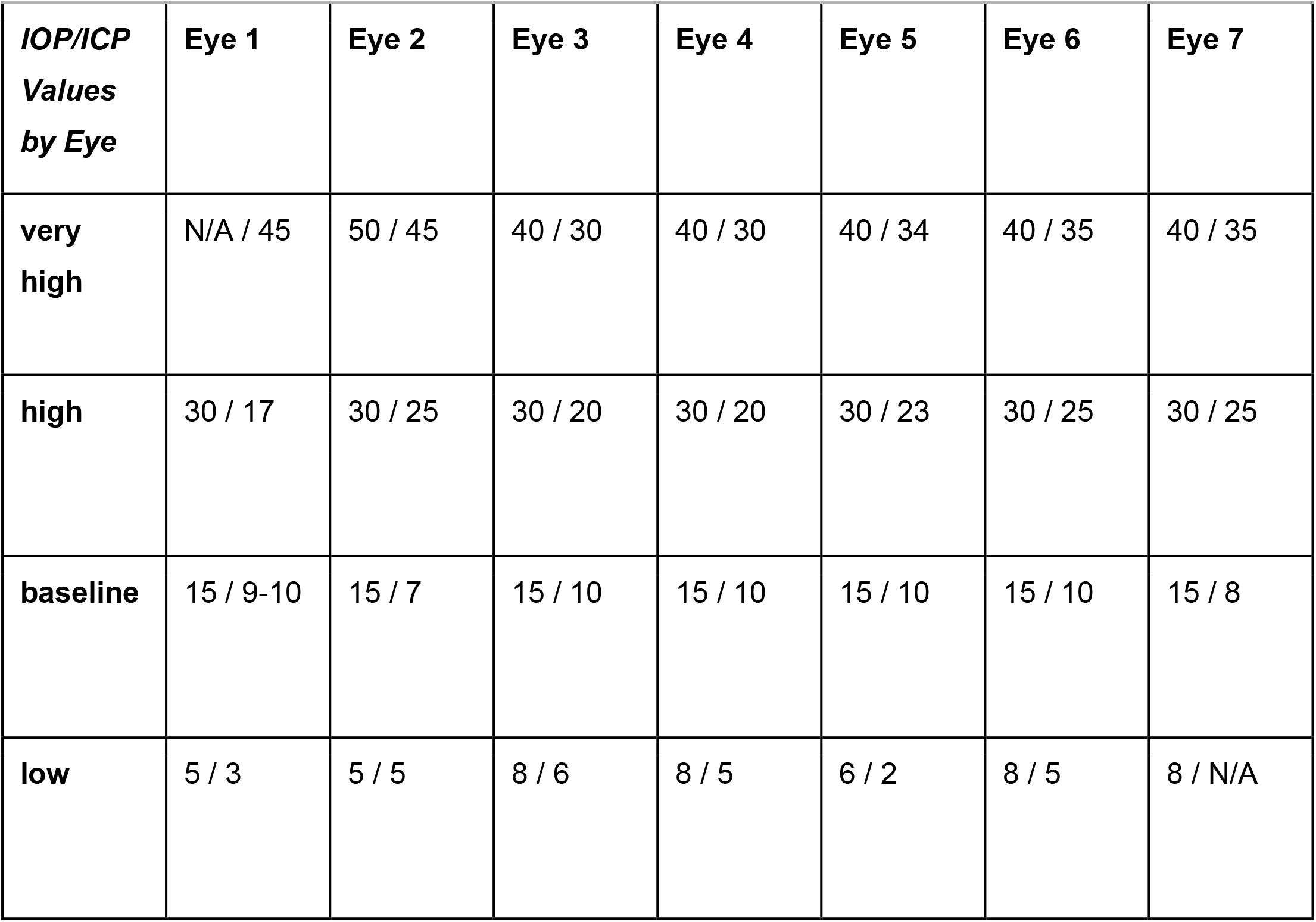
IOP and ICP values for low, baseline, high, and very high pressure groups for each eye.

